# Langerhans Cell-targeted Protein Delivery Enhances Antigen-Specific Cellular Immune Response

**DOI:** 10.1101/2025.05.05.652195

**Authors:** Ramona Rica, Klara Klein, Litty Johnson, Gabriele Carta, Mirza Sarcevic, Freyja Langer, Christoph Rademacher, Robert Wawrzinek, Federica Quattrone, Florian Sparber

## Abstract

Targeted antigen delivery to immune cells, particularly dendritic cells, has emerged as a promising strategy to enhance therapeutic efficacy of vaccines, while minimizing adverse effects associated with conventional immunization. In this study, we use our previously described small glycomimetic molecule that selectively recognizes the Langerhans cell (LC)-specific surface receptor Langerin and demonstrate specific delivery of protein antigens to these specialized dendritic cells. Our results show that Langerin-mediated antigen delivery significantly enhances the immune response *in vivo*, resulting in increased expansion and activation of antigen-specific T cells, compared to immunization with unmodified antigen. We demonstrate the feasibility of our LC-targeted platform for immune cell-specific immunization with protein antigen and underscore the potential of LCs as an access point for next-generation vaccines and immunotherapies.

## Introduction

Directing antigens actively and specifically to dendritic cells (DCs) represents a promising strategy to generate an efficient immune response with minimal use of therapeutic material. Consequently, achieving this *in vivo* is considered a major step towards the development of novel and highly efficient vaccines and immunotherapies [1]–[3].

The human skin is increasingly recognized as a valuable vaccination site, due to its dense network of DCs [4]–[6]. Amongst the different skin DC subsets, Langerhans cells (LCs) became an interesting access point for targeted immunization [5], [7]. These epidermis-residing, antigen-presenting cells can mount immune responses against invading pathogens, as well as maintain immune tolerance during tissue homeostasis. LCs use Toll-like receptors (TLRs) and C-type lectin receptors (CLRs) (such as Langerin/CD207) to recognize and engulf pathogens [8]–[10]. This process promotes LC activation and migration towards the lymph nodes, during which LC maturation and antigen processing occurs [11]. LCs have been described to orchestrate virtually all branches of immune responses by presenting antigens to CD4^+^ and CD8^+^ T cells, as well as contributing to humoral immune responses [12]–[17]. Furthermore, specific delivery of antibody-antigen complexes to LCs have been shown to enhance T cell responses [18], [19]. Conversely, LCs are capable of inducing regulatory T cells and maintain immune homeostasis and tolerance during steady state [20]–[22].

Previously, we reported the first small molecule glycomimetic ligand, selectively recognized by Langerin (CD207), a surface receptor expressed by LCs [23]. We named this technology “Langerhans Cell-Targeted Delivery System” (LC-TDS), which allows the functionalization of virtually any antigen modality to act as a LC-specific immune modulator. In contrast to antibody-mediated targeting approaches, which irreversibly bind the Langerin receptor, the LC-TDS mimics the physiological ligand-receptor interaction. We speculate that this interaction allows internalization, subsequent endosomal antigen release, and finally recycling of the receptor back to the cell surface. This LC-specific antigen delivery has been successfully demonstrated for liposomes and recombinant protein antigen in intact human skin explants [23]–[26].

In this study, the LC-TDS is used to directly and efficiently deliver protein antigens to LCs in mice, to demonstrate its efficacy to elicit productive immune responses. Immunization with LC-TDS-functionalized protein antigen, resulted in enhanced activation and expansion of antigen-specific T cells *in vivo*, compared to non-functionalized protein. This was achieved by intradermal injection as well as topical antigen deposition. Our data indicate that even minute amounts of functionalized protein is presented by LCs to cognate T cells, demonstrating the technology’s capability to promote significant antigen dose-sparing. Together, we showcase the potential of using the LC-TDS to target protein antigens directly towards LCs and thus harness the unique properties of these cells for novel immunization approaches.

## Material and Methods

### Mice

Transgenic mice expressing the human Langerin and diphtheria toxin receptor on LCs (MGI: J:113309) [27] were imported and used under the license agreement with the University of Minnesota. OT-II transgenic mice (MGI: J:87876) [28] and CD45.1 congenic mice (MGI: J:8080) [29] were purchased from The Jackson Laboratory and interbred to generate congenic OT-II/CD45.1 mice. Experimental mice were 8-12 weeks of age and were maintained in the Preclinical Facility (PCF) at the Institute of Science and Technology Austria (ISTA). Animal experiments were approved by the Austrian Federal Ministry for Education, Science and Research (animal protocol number: BMBWF-2022-0-704.335, 2024-0.025.331, 2023-0-930.309). Animal husbandry and experiments were performed in compliance with national laws and according to the FELASA (Federation of European Laboratory Animal Science Association) guidelines.

### Genotyping

Sequence of primers used for genotyping experimental animals:

LangWT fw: 5’– GAATGACAGATCTGGCCTGAGCTCG – 3’

LangWT rev: 5’– GCCAAGGCTTCTGAGAAGGATCAGG – 3’

DTR-1: 5’– GCCACCATGAAGCTGCTGCCG – 3’

DTR-2: 5’– TCAGTGGGAATTAGTCATGCC – 3’

OT-II fw: 5’– GCTGCTGCACAGACCTACT – 3’

OT-II rev: 5’– CAGCTCACCTAACACGAGGA – 3’

### Formulation of LC-TDS-functionalized protein

Green Fluorescent Protein (GFP) was generated in *E. coli* and kindly provided by Prof. Christoph Rademacher. High-quality Ovalbumin EndoFit (OVA) protein was purchased from InvivoGen. Alexa Fluor 647 conjugated Ovalbumin (OVA-AF647) protein was purchased from Sigma. Proteins containing salts and sugars underwent dialysis as a preliminary purification step prior to ligand coupling to ensure removal of interfering substances. For protein-ligand coupling, the protein was dissolved in phosphate-buffered saline (PBS) pH 8 (Gibco) in a glass vial. The linker-ligand solution was prepared by dissolving the linker-ligand in anhydrous N,N-Dimethylformamid (DMF) (Sigma). The detailed procedure for the linker-ligand conjugation was previously described [25]. Under gentle agitation on a rocker shaker, the linker-ligand solution was added to the protein, and the mixture was incubated at room temperature (RT) for 17 hours to complete the reaction. Following the reaction, the protein-ligand conjugate was purified by dialysis twice against PBS at 4°C using Slide-A-Lyzer Dialysis Cassettes (7K MWCO, Thermo Fisher Scientific). Dialyzed samples were carefully collected from the cassette using a syringe to avoid sample loss. Protein concentration was determined using NanoPhotometer NP80 (Implen) and Bicinchoninic Acid (BCA) assay. The coupling efficiency was assessed by nuclear magnetic resonance (NMR) spectroscopy.

### Intradermal immunization

Animals were anesthetized by intraperitoneal (i.p.) injection of Ketamine (final: 90 mg/kg) and Xylazine (final: 7.5 mg/kg). Protein formulations were diluted in PBS and a total of 25 µl/ear was injected intradermally (i.d.) into the ear pinnae using an insulin syringe (Omnican 30G, B. Braun).

### Topical application of protein antigen

For topical application, animals were anesthetized as previously described. When indicated, mouse ears were tape-stripped 5x using Transpore Surgical Tape (3M) before applying the protein antigen. Protein formulations were diluted 1:1 in Ultrasicc (prepared at the Pharmacy) and a total of 15 µl/ear was applied onto the ears and spread with a pipette tip.

### Generation of epidermal cell (EC) suspensions from ear and body skin

Epidermal cell (EC) suspension from ear and body skin was prepared as previously described [30]. Briefly, the fur was removed by plucking. After body skin removal, subcutaneous fat was removed by scraping with a scalpel blade, subsequently incubating smaller skin pieces with the dermal side down in 0.6% Trypsin (Sigma) solution. Ears were cut at the base, split into ventral and dorsal halves and the latter ones were incubated dermal side down in 0.6% Trypsin solution. After 20-30 min incubation at 37°C, skin pieces and ears were transferred directly into fetal bovine serum (FBS) (Cytiva) dermal side down. Using forceps, epidermis was peeled off from dermis and transferred into complete medium (RPMI1640 (Gibco) supplemented with 10% FBS, 1% Penicillin/Streptomycin (Sigma) and L-Glutamine (Sigma)). Epidermal skin pieces were further disrupted using forceps. Medium together with epidermal skin and FBS was collected and incubated for another 30 min at 37°C in the water bath (shaking) for further disruption. Single cell suspension was generated by straining the suspension and mashing the remaining epidermal skin pieces through a 100 µm cell strainer (Sarstedt).

### Lymph node and spleen cell suspension

Single cell suspension was generated by mashing organs through a 70 µm cell strainer (Sarstedt). For erythrocyte lysis of spleen samples, cell pellet was resuspended in Red Blood Cell Lysis Buffer (Roche) and incubated for 3 min at RT. After washing, cells were used for staining.

### OT-II T cell isolation and adoptive T cell transfer

Naïve OT-II CD4^+^ T cells were isolated using the naïve CD4^+^ T Cell Isolation Kit, mouse (Miltenyi Biotec) according to the manufacturer’s protocol. In summary, all peripheral lymph nodes (except mesenteric lymph nodes) and spleen were isolated from OT-II/CD45.1 mice and pooled. After erythrocyte lysis, OT-II T cells were enriched using the negative selection kit and T cells were labelled with Carboxyfluorescein succinimidyl ester (CFSE, 1:2000, BioLegend) for 20 min at 37° C. After washing, cells were resuspended in either complete medium for the EC:OT-II co-culture or in PBS for adoptive T cell transfer. A total of 1×10^6^ cells were transferred intravenously (i.v.) into mice.

### Diphtheria toxin injection

For depletion of Langerhans cells (LCs) *in vivo*, huLang animals (expressing the diphtheria toxin receptor under the control of the human Langerin promoter) were injected i.p. with 150 ng of diphtheria toxin (DT) (Sigma) four days prior to the adoptive T cell transfer.

### Co-culture of EC suspension and OT-II T cells

Co-cultures were set up in two different settings. EC suspension generated from whole body skin of naïve huLang mice was co-cultured together with naïve CFSE-labelled OT-II T cells in the presence of t- or nt-OVA protein formulations (1 µg/ml, 10 µg/ml and 100 µg/ml). When mice were treated intradermally or topically with t- or nt-OVA protein formulations (5 µg/ear), EC suspension from the treated ears was generated 3 hours post application and subsequently incubated with naïve CSFE-labelled OT-II T cells. Co-cultures were prepared at a ratio of 20:1 (EC:OT-II) and incubated for 3 days at 37°C and 5% CO_2_.

### Flow cytometric analysis

After washing, cells were incubated with LIVE/DEAD™ Fixable Near IR (780) Viability Kit or LIVE/DEAD™ Fixable Yellow Dead Cell Stain Kit (1:1000; Invitrogen, Thermo Fisher Scientific) for 20 min on ice, subsequently washing again with PBS and adding the surface antibody mix for 20 min on ice. For the detection of intracellular human Langerin receptor, BD Cytofix/Cytoperm Fixation Buffer and BD Perm/Wash Buffer (BD Biosciences) were used. After extracellular staining, cells were fixed for 20 min on ice, followed by the intracellular staining for 20 min on ice. Cells were measured with a CytoFLEX Flow Cytometer S (Beckman Coulter) and analyzed by FlowJo v10.6.1 (BD Bioscience).

### Flow cytometry antibodies

Following antibodies were used for this study: anti-mouse I-A/I-E (MHC-II) (M5/114.15.2), anti-mouse CD69 (H1.2F3), anti-mouse CD44 (IM7), anti-mouse CD25 (PC61), anti-mouse CD4 (GK.1), anti-mouse CD8a (53-6.7), anti-mouse CD45.1 (A20) were purchased from BioLegend; anti-human CD207 (Langerin) (MB22-9F5) was purchased from Miltenyi Biotec.

### Statistical analysis

Statistical analysis was performed using Prism 10 (GraphPad). One-way ANOVA followed by Tukey’s multiple-comparison test was used for the comparison of more than 2 groups. The *p*-values are defined as follows: *p <0.05; **p <0.01; ***p <0.001; ****p <0.0001; n.s., not significant.

## Results

### Selective uptake of LC-TDS-functionalized protein antigen by Langerhans cells *in vitro*

To investigate the potential of the LC-TDS-functionalized protein to elicit antigen-specific immune responses, we conducted a series of *in vitro* and *in vivo* experiments. We refer to protein antigens conjugated with the LC-TDS as “targeted” (t-)proteins, while unmodified proteins, serving as controls, are referred to as “non-targeted” (nt-)proteins. Considering that the LC-TDS specifically targets the human Langerin receptor [23]–[25], we utilized transgenic mice expressing human Langerin on the surface of LCs (referred to as huLang mice [27]). To confirm that t-proteins are specifically delivered to LCs, we incubated epidermal cell (EC) suspension, generated from huLang mice, with targeted- or non-targeted green fluorescent protein (t- or nt-GFP) for 2 hours and subsequently analyzed the protein uptake by flow cytometry (**Fig. 1A**). EC suspension primarily consists of keratinocytes, alongside LCs, the sole DC subset in this tissue, which can be distinguished by the expression of MHC-II (**Fig. 1B**). Using two protein concentrations (5 and 50 µg/ml), we observed that neither t-nor nt-GFP induced cell death in LCs or keratinocytes (**Fig. 1C**). Further examination revealed that only LCs but not keratinocytes, selectively internalized t-GFP. LCs did not internalize nt-GFP during the 2-hour incubation (**Fig. 1D** and **E**). Keratinocytes appeared to take up t- and nt-GFP just to a minor degree (∼1%) and only at high protein concentration. In contrast, t-GFP is greatly internalized by LCs (∼80-90%) at low and high protein concentration (**Fig. 1D** and **E**). Interestingly, the uptake of t-GFP by LCs *in vitro* reached saturation already at the lower concentration of 5 µg/ml (**Fig. 1E**). This data indicates that the LC-TDS enables selective and non-toxic delivery of protein antigens specifically to LCs, with efficient uptake reaching saturation already at low protein concentrations.

**Fig. 1:**
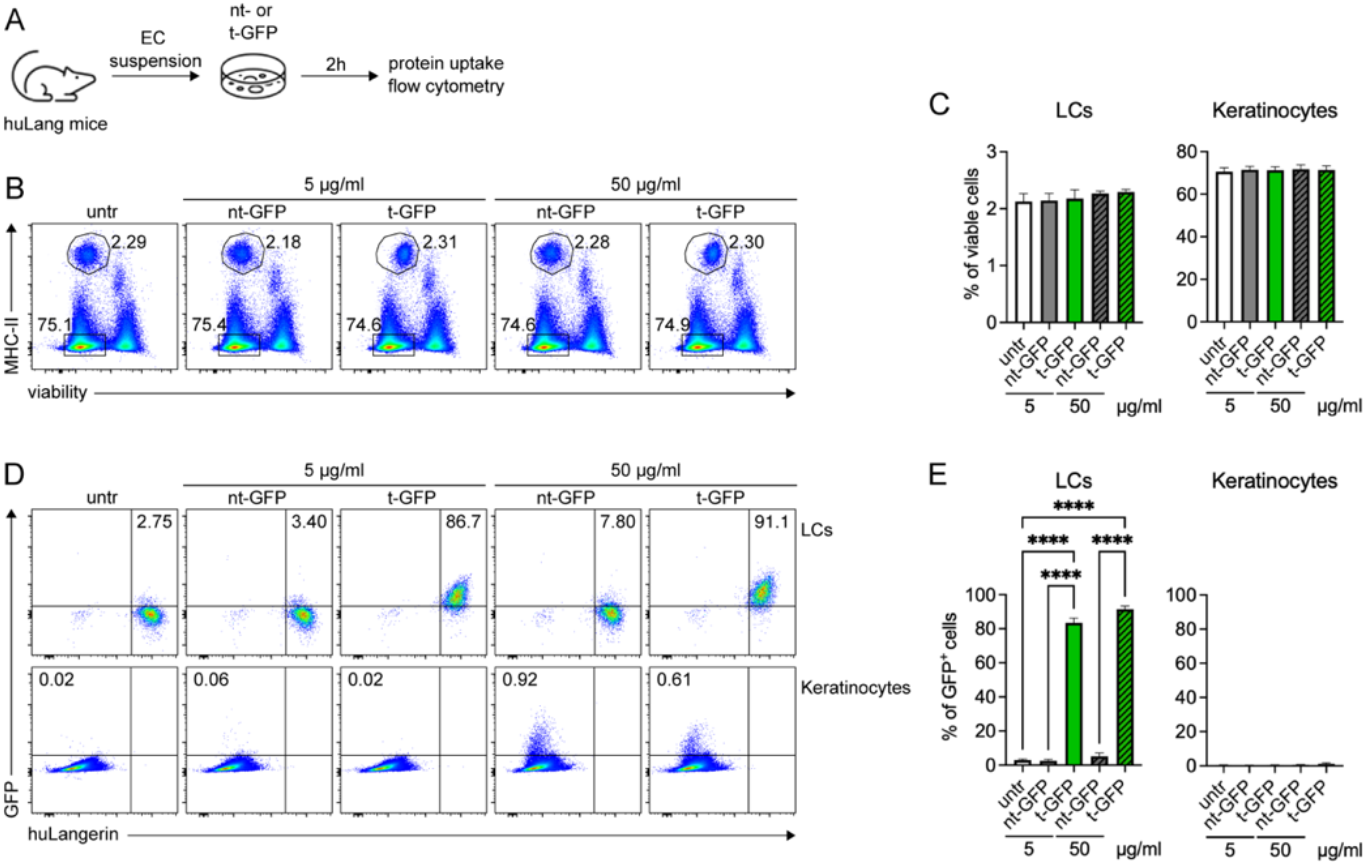
Selective uptake of LC-TDS-functionalized protein antigen by LCs *in vitro*. **(A)** Scheme of experimental setup. Epidermal cell (EC) suspension was generated from huLang mice and incubated with t- or nt-GFP at two concentrations (5 and 50 μg/mL) for 2 hours. Protein uptake was determined by flow cytometry. **(B)** Representative plots show gating strategy of MHC-II^+^ LCs and MHC-II^-^ keratinocytes for all treatment conditions and untreated (untr) control. **(C)** Graphs show the percentages of viable MHC-II^+^ LCs (left) or MHC-II^-^ keratinocytes (right). **(D)** Representative plots show the expression of human Langerin (huLang) and the uptake of GFP by LCs (upper panel) or keratinocytes (lower panel). **(E)** Graphs show the percentages of huLangerin^+^GFP^+^ LCs (left) or huLangerin^-^GFP^+^ keratinocytes (right). **(B** and **D)** Numbers in the plots represent the percentage of cells within the indicated regions. All graphs show pooled data from four independent experiments with 5-6 replicates/group. Statistics were calculated using a one-way ANOVA, \*\*\*\**p* < 0.0001.

### Langerhans cell-specific uptake of LC-TDS-functionalized protein antigen *in vivo*

Next, we aimed to investigate whether the selective uptake of t-GFP protein by LCs could also be observed after antigen administration *in vivo*. Furthermore, we tested whether our LC-targeting technology is broadly applicable for the delivery of various types of protein antigens to LCs in the skin. To this end, in addition to GFP, we formulated nt- and t-Ovalbumin (OVA), fluorescently labelled with Alexa Fluor 647 (OVA-AF647), and injected the proteins intradermally (i.d.) into the ear pinnae of huLang mice. Protein uptake was analyzed 3 hours after antigen application in EC suspension generated from the injection site (**Fig. 2A**).

**Fig. 2:**
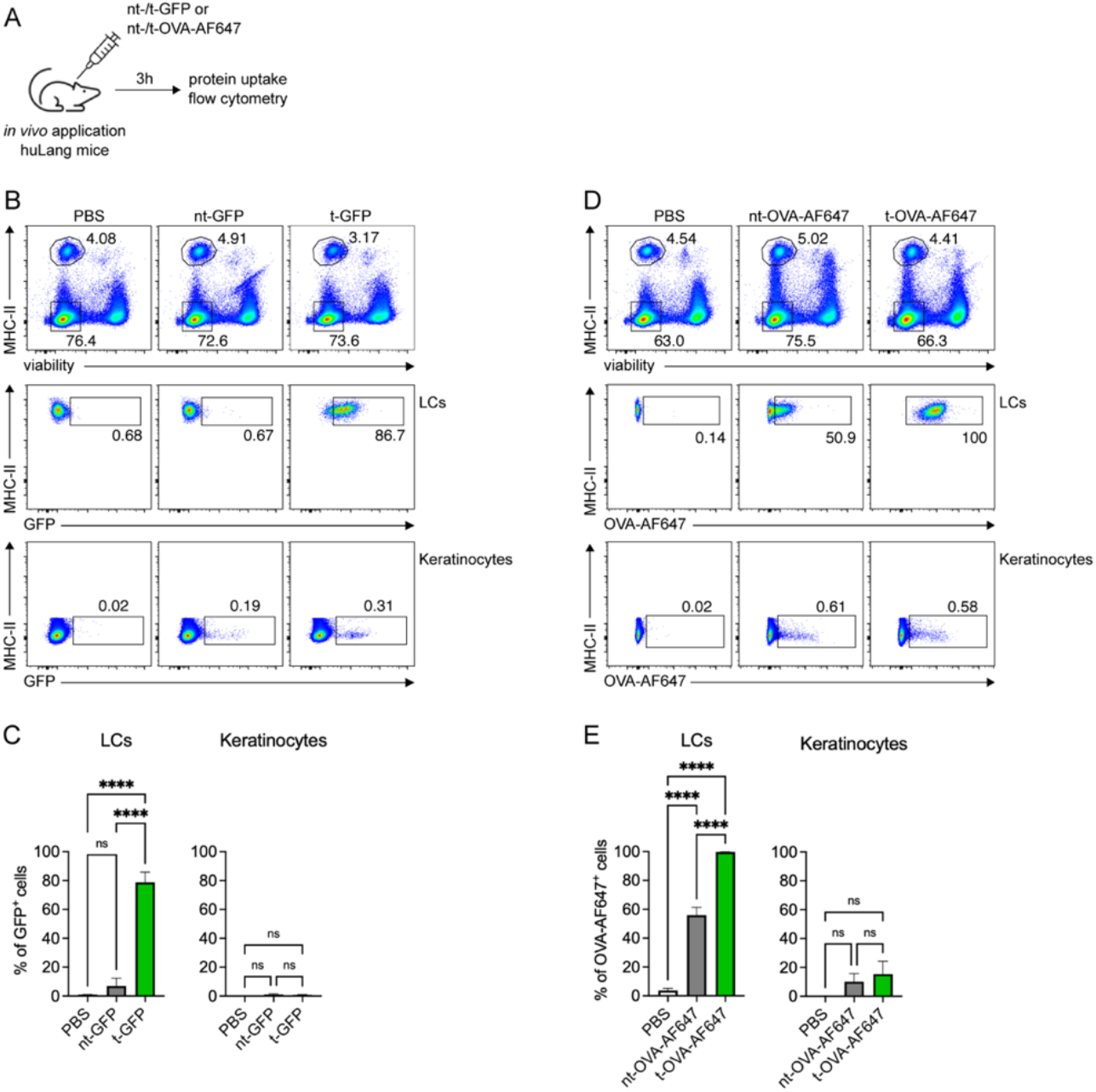
Langerhans cell-specific uptake of LC-TDS-functionalized protein antigen *in vivo*. **(A)** Scheme of experimental setup. HuLang mice were injected i.d. with 5 µg/ear of GFP or OVA-AF647 formulations and protein uptake was determined by flow cytometry 3 hours later. **(B** and **D)** Representative plots show gating strategy (upper panel) and uptake of protein antigen (middle and lower panel) by LCs (MHC-II^+^) and keratinocytes (MHC-II^-^) from EC suspension, after *in vivo* application of either **(B)** t- or nt-GFP or **(D)** t- and nt-OVA-AF647. **(C** and **E)** Graphs show the percentage of **(C)** GFP^+^ or **(E)** OVA-AF647^+^ cells out of MHC-II^+^ LCs (left) or MHC-II^-^ keratinocytes (right). All graphs show pooled data from two independent experiments with 4-6 replicates/group. Statistics were calculated using a one-way ANOVA, \*\*\*\**p* < 0.0001, n.s. p ≥ 0.05.

Similar to the *in vitro* setup, we distinguished between LCs and keratinocytes, based on MHC-II expression (**Fig. 2B** and **D**). Examination of GFP fluorescence in these cell populations confirmed that only the t-GFP, but not nt-GFP, was efficiently internalized by LCs (∼80%). Keratinocytes showed minimal detectable uptake of both nt- and t-GFP formulations (∼0.3%) (**Fig. 2B** and **C**). Interestingly, internalization of OVA-AF647 formulations appeared to behave differently from GFP. While we observed again a very strong uptake of t-OVA-AF647 by LCs (∼100%), nt-OVA-AF647 was also taken up by LCs, though to a significant lesser extent (∼50%) (**Fig. 2D** and **E**). Similar to the internalization of GFP protein, keratinocytes showed minimal uptake of both nt- and t-OVA-AF647 protein (∼5%). Overall, the functionalization of protein antigens with the LC-TDS significantly enhances LC-specific uptake *in vivo* upon i.d. administration.

### Enhanced antigen presentation of LC-TDS-functionalized protein by Langerhans cells

So far, we have demonstrated that LC-TDS-functionalized protein antigen is selectively taken up by LCs and is internalized to a significantly greater extent than its corresponding nt-formulation (**Fig. 2D** and **E**). To determine whether this uptake also correlates with enhanced antigen presentation to T cells, we focused on antigen-presentation to CD4^+^ T cells, as exogenous protein antigens will mainly be presented to MHC-II-restricted T cells. Co-culturing cognate CD4^+^ T cells with EC suspension allowed us to investigate LC-specific antigen presentation, as LCs are the only MHC-II positive cells within the epidermis. Herein, we first co-cultured EC suspension, generated from naïve huLang mice, with naïve CFSE-labelled OVA-specific CD4^+^ (OT-II) T cells, in the presence of increasing concentrations of either t- or nt-OVA protein. After 3 days of co-culture, we investigated T cell proliferation by CFSE-dilution (**Supp. Fig. 1A**). The analysis showed, that even low protein concentrations of t-OVA (1 µg/ml) led to a significantly increased T cell expansion, compared to nt-OVA (∼53% vs. ∼4%, respectively) (**Supp. Fig. 1B** and **C**). While T cell proliferation induced by LCs in the presence of nt-OVA was dose-dependent, even at the highest protein concentration (100 µg/ml), nt-OVA was significantly outperformed by t-OVA (∼36% vs. ∼54%, respectively) (**Supp. Fig. 1C**). Notably, LC-mediated T cell proliferation induced by incubation\ with t-OVA reached saturation even at the lowest concentration (1 µg/ml), exceeding the proliferation induced by the highest concentration of nt-OVA (100 µg/ml). This indicates a substantial dose-sparing effect of LC-TDS-mediated antigen delivery.

**Supp. Fig. 1:**
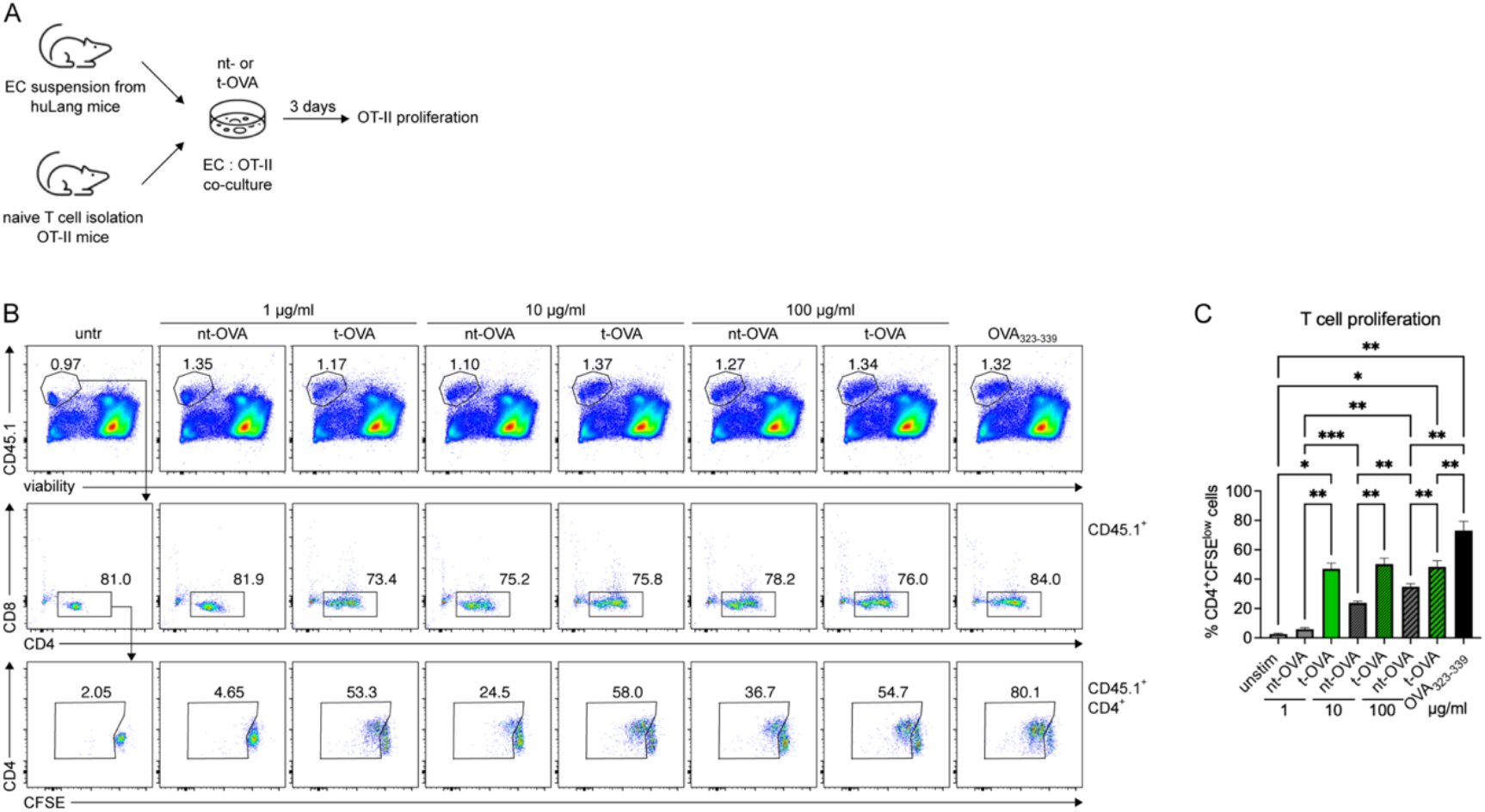
LC-specific protein delivery results in a pronounced antigen dose sparing *in vitro*. **(A)** Scheme of experimental setup. EC suspension generated from naïve huLang mice was co-cultured for 3 days with naïve CFSE-labelled OT-II T cells in the presence of increasing concentrations of t- or nt-OVA (1, 10 or 100 μg/ml). **(B)** Representative plots show the gating strategy for CD45.1^+^CD4^+^ OT-II T cells (upper and middle panel) and corresponding T cell proliferation by CFSE dilution (lower panel). **(C)** Graph shows the percentages of proliferating CD4^+^CFSE^low^ OT-II T cells after co-culture. All graphs show pooled data from three independent experiments with 5-10 replicates/group. Statistics were calculated using a one-way ANOVA, \**p* < 0.05, \*\**p* < 0.01, \*\*\**p* < 0.001.

We next sought to investigate, whether LCs also effectively process and present antigen to T cells upon *in vivo* antigen administration. We conducted an experimental series in which we applied t- or nt-OVA (5 µg) by two methods. Additional to the already described i.d. injection into the ear pinnae, we also tested topical application onto the dorsal ear skin. For this, the skin barrier was disrupted by tape-stripping prior to antigen application. Three hours following the administration, EC suspension was generated from treated ears and co-cultured with naïve CFSE-labelled OT-II T cells (**Fig. 3A**). Analysis after three days of co-culture showed that only t-OVA effectively promoted T cell responses, as evidenced by strong T cell proliferation upon co-culture with EC suspension from mice treated i.d. or topically with t-OVA (**Fig. 3B**). LC-mediated T cell proliferation reached ∼46% after i.d. injection (**Fig. 3C**) and ∼20% after topical application of t-OVA (**Fig. 3D**). Notably, application of nt-OVA *in vivo* induced significantly lower T cell proliferation *in vitro* (∼3%), irrespective of the administration route (**Fig. 3C** and **D**). The induction of proliferation positively correlated with the expression levels of early CD4^+^ T cell activation markers. CD69, CD44 and CD25 were significantly upregulated by T cells upon co-culture with EC suspension from mice treated either i.d. (**Fig. 3E** and **F**) or topically (**Fig. 3G** and **H**) with t-OVA compared to nt-OVA.

**Fig. 3:**
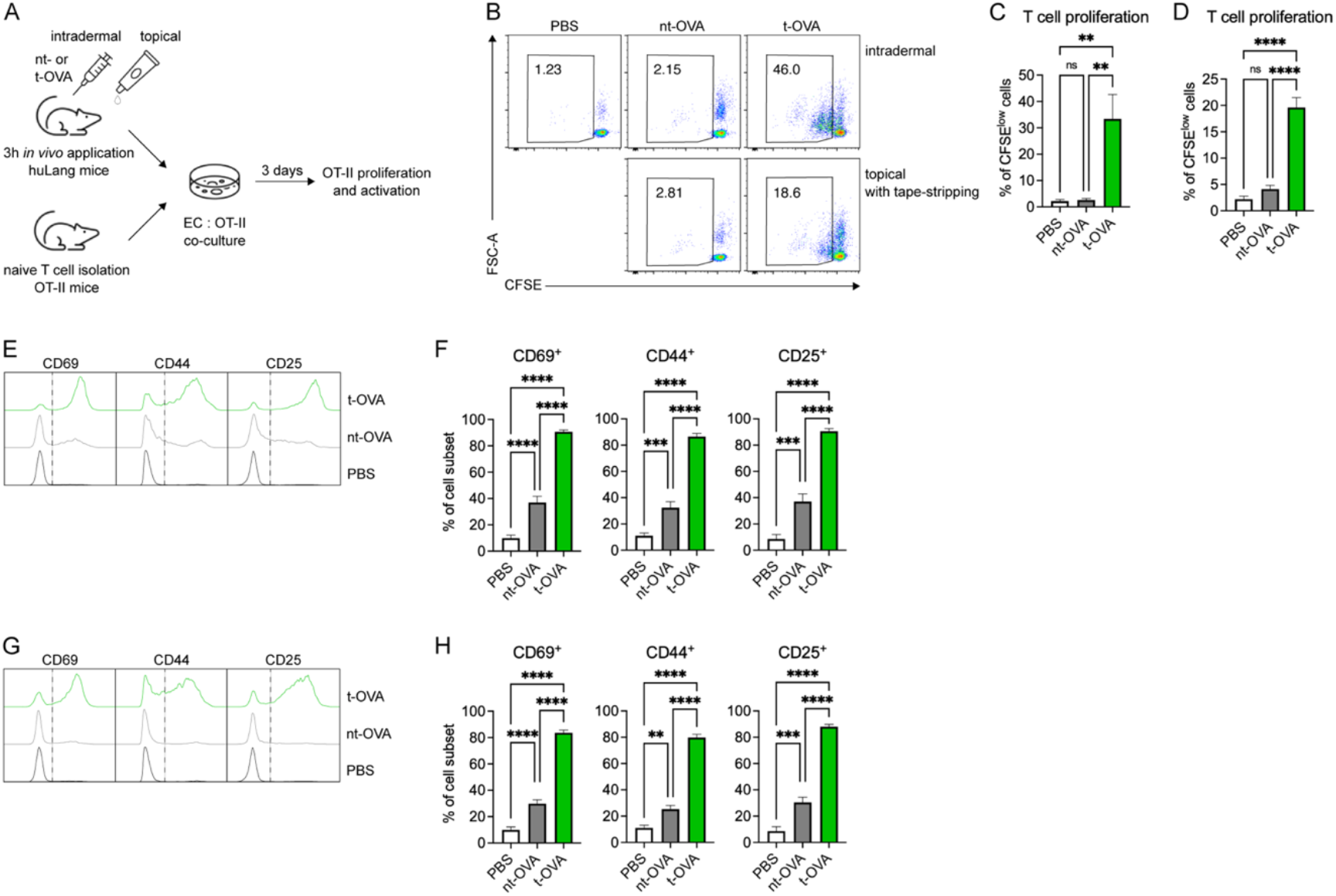
LC-targeted protein delivery *in vivo* induces a robust T cell activation *in vitro*. **(A)** Scheme of experimental setup. HuLang mice were either i.d. or topically treated with 5 µg/ear of t- or nt-OVA. Three hours following the administration, EC suspension was generated from the treated ears and co-cultured with naïve CFSE-labeled OT-II T cells for 3 days *in vitro*. T cell activation and proliferation was assessed by flow cytometry. **(B)** Representative plots show T cell proliferation based on CFSE dilution, after co-culture with EC suspension from i.d. (upper panel) or topically (tape-stripped; lower panel) treated mice. **(C** and **D)** Graphs show the percentages of proliferating CD4^+^CFSE^low^ T cells after co-culture with EC suspension from **(C)** i.d. or **(D)** topically treated mice. **(E – H)** Histograms and corresponding graphs showing expression of CD69, CD44 and CD25 by CD4^+^ OT-II cells, co-cultured with EC suspension from **(E** and **F)** i.d. or **(G** and **H)** topically treated mice. All graphs show pooled data from two independent experiments with 5-6 replicates/group. Statistics were calculated using a one-way ANOVA, \*\**p* < 0.01, \*\*\**p* < 0.001, \*\*\*\**p* < 0.0001, n.s. p ≥ 0.05.

Furthermore, in a similar experiment LCs, efficiently captured t-OVA protein and promote superior T cell response, even when applied topically onto intact skin, without tape-stripping (**Supp. Fig. 2 A – D)**. Taken together, these findings indicate that immunization with LC-TDS-functionalized protein *in vivo* results in superior T cell activation *in vitro*, even at low protein concentrations.

**Supp. Fig. 2:**
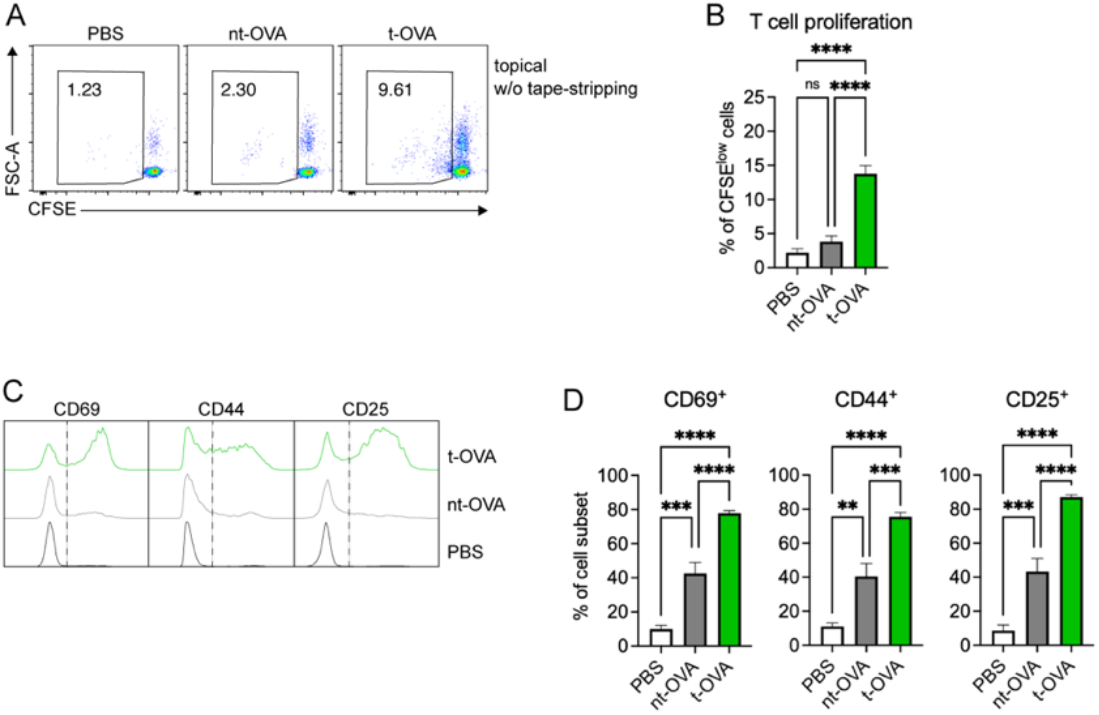
Topical administration of LC-TDS-functionalized protein onto intact skin results in enhanced T cell activation *in vitro*. **(A)** Representative plots show OT-II T cell proliferation based on CFSE dilution after co-culture with EC suspension from mice treated topically with 5 µg/ear of t- or nt-OVA (without tape stripping). **(B)** Graphs show percentages of proliferating CD4^+^CFSE^low^ OT-II T cells after co-culture. **(C** and **D)** Histograms and corresponding graphs showing expression of CD69, CD44 and CD25 by CD45.1^+^CD4^+^ OT-II T cells, co-cultured with EC suspension from topically treated mice. All graphs show pooled data from two independent experiments with 5-6 replicates/group. Statistics were calculated a using one-way ANOVA, \*\**p* < 0.01, \*\*\**p* < 0.001, \*\*\*\**p* < 0.0001, n.s. p ≥ 0.05.

### Enhanced *in vivo* T cell response upon immunization with LC-TDS-functionalized protein antigen

Our data suggests that specific delivery of low amounts of protein antigen to LCs *in vivo*, by means of the LC-TDS, result in a superior T cell activation. Lastly, we explored whether immunization with t-OVA leads to a similar enhanced immune response *in vivo*, using adoptive OT-II T cell transfer. To confirm that the LC-TDS-mediated enhancement of the T cell proliferation and activation is dependent on the presence of LCs and their expression of the human Langerin receptor, we also immunized huLang mice, in which LCs were specifically depleted by diphtheria toxin (DT) administration (due to the LC-specific expression of the diphtheria toxin receptor in these mice), as well as murine Langerin-expressing wild-type (WT) mice, with t-OVA.

Naïve CFSE-labelled OT-II T cells were adoptively transferred one day prior to the i.d. immunization with the OVA formulations. T cell proliferation was assessed seven days post-immunization (**Fig. 4A**). Significant expansion in absolute numbers and frequencies of total cells and CD45.1^+^CD4^+^ OT-II T cells was detected in the draining lymph nodes (LN) from LC-sufficient huLang mice immunized with t-OVA. LC-depleted huLang (huLang + DT) and WT mice immunized with t-OVA, as well as huLang mice treated with nt-OVA showed significantly less OT-II T cell expansion (**Fig. 4B-D**). Consistently, we observed a significantly higher frequency of proliferating CFSE^low^ OT-II T cells in huLang mice immunized with t-OVA, compared to control animals (**Fig. 4E**).

**Fig. 4:**
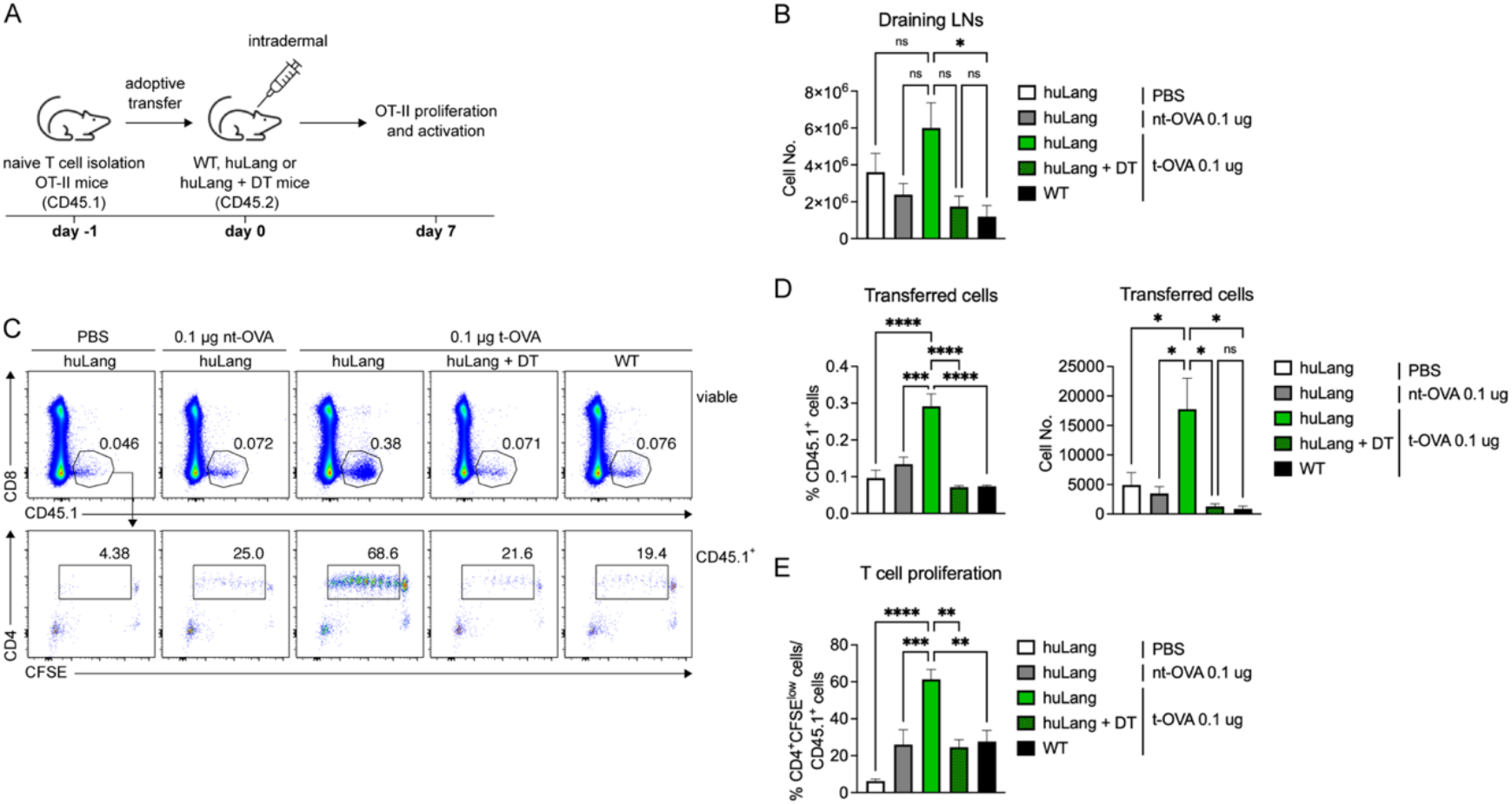
Enhanced *in vivo* T cell response upon immunization with LC-TDS-functionalized protein antigen. **(A)** Scheme of experimental plan. Naïve CFSE-labelled OT-II T cells were adoptively transferred one day prior to the immunization with 0.1 µg of t- or nt-OVA. OT-II T cell expansion and proliferation was assessed on day 7 post immunization (p.i.) by flow cytometry. **(B)** Graph shows total cell numbers in draining lymph nodes (LNs) 7 days p.i. **(C)** Representative plots showing gating strategy (upper panel) and corresponding proliferation (lower panel) of CD4^+^CFSE^low^ OT-II T cells within the LNs 7 days p.i. **(D)** Graph shows the percentages (left) and absolute cell numbers (right) of transferred CD45.1^+^CD4^+^ OT-II T cells. **(E)** Graph shows the percentage of proliferating CD45.1^+^CD4^+^CFSE^low^ OT-II T cells. All graphs show pooled data from two independent experiments with 4-6 replicates/group. Statistics were calculated using a one-way ANOVA, \**p* < 0.05, \*\**p* < 0.01, \*\*\**p* < 0.001, \*\*\*\**p* < 0.0001, n.s. p ≥ 0.05.

While we could not observe substantial differences in the absolute cell numbers of splenic CD45.1^+^CD4^+^ OT-II T cells between the different treatment groups, we again detected a significant increase in proliferating CD4^+^CFSE^low^ OT-II T cells in the spleen of huLang mice immunized with t-OVA protein (∼15%), compared to control animals (**Supp. Fig 3A** – **D**).

**Supp. Fig. 3:**
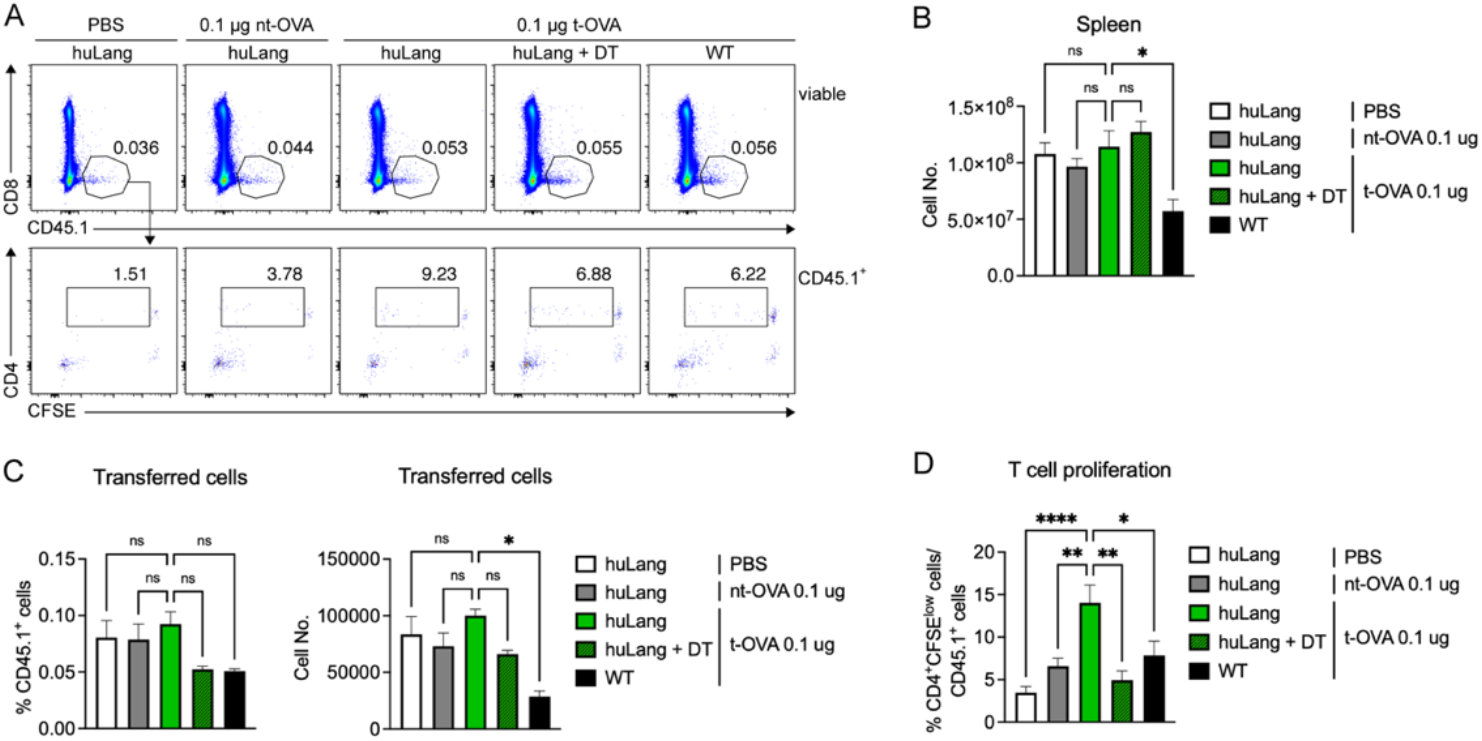
Increased proliferation of splenic antigen-specific T cells upon immunization with LC-TDS-functionalized protein. **(A)** Representative plots show percentage of transferred OT-II T cells (upper panel) and proliferating CD4^+^CFSE^low^ OT-II T cells (lower panel), 7 days post immunization within the spleen. **(B)** Graph shows absolute cell numbers of solenocytes from immunized mice. **(C)** Graphs show the percentages (left) and absolute cell numbers (right) of transferred CD45.1^+^CD4^+^ OT-II T cells within the spleen. **(D)** Graph shows the percentages of proliferating CD4^+^CFSE^low^ OT-II T cells. All graphs show pooled data from two independent experiments with 4-8 replicates/group. Statistics were calculated using a one-way ANOVA, \**p* < 0.05, \*\**p* < 0.01, \*\*\*\**p* < 0.0001, n.s. p ≥ 0.05.

This data confirms that the immunization with LC-TDS-functionalized protein induces a superior cellular immune response *in vivo*, which is dependent on the presence of LCs and their expression of human Langerin receptor.

## Discussion

The precise delivery of antigens to DCs *in vivo* offers significant advantages over conventional immunization strategies by enhancing antigen uptake efficiency, while reducing off-target effects and minimizing potential reactogenicity as well as adverse effects for the host [31]. LCs, the sole DC subset in the epidermis, are particularly suited for skin-targeted immunization approaches. Our study builds on previous research about LC-specific antigen delivery using a small glycomimetic ligand, and now demonstrates the functionality of the LC-TDS platform to evoke a potent cellular immune response *in vivo* [23]–[25].

We showed that LC-TDS-functionalized protein antigens enable specific and more efficient uptake by LCs *in vitro* compared to non-targeted antigens. This subsequently led to superior antigen-specific T cell proliferation, even at low antigen concentrations of 1 µg/ml, and surpassed the proliferation observed with 100 µg/ml of non-targeted antigen. Consequently, LC-specific antigen targeting can reduce the required antigen dose for an adequate T cell response by at least two-log fold. Such dose-sparing, in combination with the minimally invasive immunization through the skin, represents a transformative prospect for the development of novel vaccines with higher efficacy, while minimizing possible adverse effects. Additionally, the ability to reduce antigen dose while maintaining immunogenicity also reduces the overall cost of production and conserves the antigenic material thereby making vaccines more accessible. This is particularly beneficial in resource limited settings where manufacturing constraints and distribution challenges often limit widespread immunization [32]. These benefits are relevant for various therapeutic interventions including antiviral vaccines, DC-based cancer immunotherapies and antigen-specific immunotherapies (AITs) for allergies or autoimmune diseases.

Consistently, our findings reveal that immunization with LC-TDS-functionalized protein antigens led to LC-specific uptake *in situ* and enhanced T cell activation and proliferation *in vitro* and *in vivo*. This superior T cell response depended on the presence of LCs and their expression of the human Langerin receptor, as depletion or absence of either reduced the reaction to levels comparable to conventional non-targeted immunization. Moreover, the elevated cellular response was evident in intradermal and topical immunization when using targeted antigen, highlighting the administration flexibility of LC-TDS-targeted antigen delivery. Because LCs are known to efficiently capture topical antigens by extending their dendrites toward the skin surface [33], this steady-state antigen sampling is believed to be essential for inducing and maintaining immune tolerance—a mechanism currently being leveraged in the development of novel AIT for peanut allergy [34]. Our results demonstrate that topically applied LC-TDS-functionalized antigen enhances LC-mediated antigen uptake and presentation. Thus, the LC-TDS could help refine epicutaneous AIT (EPIT) for allergies and autoimmune diseases in the future.

Despite these promising results, we see certain limitations and challenges in our study, which need to be addressed: while the targeted antigen resulted in overall enhanced uptake by LCs, the extent of this difference compared to the non-targeted antigen varied depending on the model antigen used. The observed differences in antigen uptake between targeted/non-targeted GFP and targeted/non-targeted OVA-647 may be due to antigen glycosylation (GFP is non-glycosylated, whereas OVA is). The glycosylation status of an antigen is a determining factor for receptor-mediated uptake and hence targeted antigen-delivery to DCs [31]. Future experiments using antigens with different glycosylation pattern may help to fully understand the impact of antigen glycosylation on LC-TDS-mediated antigen delivery. LC-specific antigen delivery enhanced the activation and proliferation of OVA-specific OT-II T cells *in vivo*. We acknowledge that an adoptive T cell transfer does not fully replicate the induction of a *de novo* immune responses in a naïve host upon immunization. However, eliciting an endogenous immune response against a low-immunogenic protein often requires carefully selected adjuvants to generate a robust and measurable effect. While this study primarily aimed to investigate the overall potential and mechanism of the LC-TDS platform, further experiments involving the immunization of naïve animals will be essential to assess its efficacy in a more physiological context.

## Conclusion

Collectively, our findings substantiate the LC-TDS technology as a promising platform for developing next-generation targeted vaccines and immunotherapies. By directing antigens to LCs in the skin, this approach has the potential to enhance therapeutic efficacy while mitigating systemic side effects through dose-sparing strategies. This is promising for potential applications ranging from vaccines for infectious disease to tolerance induction in allergy and autoimmunity. Continued development and evaluation of this strategy in diverse immunological settings will be key to recognize its clinical potential.

## Acknowledgments

This project was generously supported by “*Seedfinancing”* (P2282679) of the Austrian BMDW and the BMK, handled by the Austrian Wirtschaftsservice (aws), as well as by “*Life Science Call 2022”* (FO999896442) of the Austrian Research Promotion Agency (FFG). We thank Mag. Michael Schunn from the Preclinical Facility (PCF) of the Institute of Science and Technology Austria for his continuous technical support.

## Author Contribution

Conceptualization: RR with support from FQ and FS;

Methodology: RR with support from KK, LJ, GC, MS, FL and FQ;

Formal Analysis: RR with support from KK and FQ;

Writing: Original draft: RR and FS, Review: CR, RW, KK and FQ;

Visualization: RR with support from FS;

Funding Acquisition: RW and FS;

## References

[1] R. L. Sabado, S. Balan, and N. Bhardwaj, “Dendritic cell-based immunotherapy,” Cell Res. 2017 271, vol. 27, no. 1, pp. 74–95, Dec. 2016, doi: 10.1038/cr.2016.157.

[2] I. Caminschi, E. Maraskovsky, and W. R. Heath, “Targeting Dendritic Cells in vivo for Cancer Therapy,” Front. Immunol., vol. 3, no. FEB, 2012, doi: 10.3389/FIMMU.2012.00013.

[3] Y. Pastor, N. Ghazzaui, A. Hammoudi, M. Centlivre, S. Cardinaud, and Y. Levy, “Refining the DC-targeting vaccination for preventing emerging infectious diseases,” Front. Immunol., vol. 13, p. 949779, Aug. 2022, doi: 10.3389/FIMMU.2022.949779/PDF.

[4] D. van Dinther, D. A. Stolk, R. van de Ven, Y. van Kooyk, T. D. de Gruijl, and J. M. M. den Haan, “Targeting C-type lectin receptors: a high-carbohydrate diet for dendritic cells to improve cancer vaccines,” J. Leukoc. Biol., vol. 102, no. 4, p. 1017, Oct. 2017, doi: 10.1189/JLB.5MR0217-059RR.

[5] N. Romani, V. Flacher, C. H. Tripp, F. Sparber, S. Ebner, and P. Stoitzner, “Targeting skin dendritic cells to improve intradermal vaccination,” Curr. Top. Microbiol. Immunol., vol. 351, no. 1, pp. 113–138, 2012, doi: 10.1007/82_2010_118.

[6] N. Romani, M. Thurnher, J. Idoyaga, R. M. Steinman, and V. Flacher, “Targeting of antigens to skin dendritic cells: possibilities to enhance vaccine efficacy,” Immunol.Cell Biol., vol. 88, pp. 424–430, 2010.

[7] P. Stoitzner, F. Sparber, and C. H. Tripp, “Langerhans cells as targets for immunotherapy against skin cancer,” Immunol. Cell Biol., vol. 88, no. 4, pp. 431–437, May 2010, doi: 10.1038/ICB.2010.31.

[8] C. G. Figdor, Y. Van Kooyk, and G. J. Adema, “C-type lectin receptors on dendritic cells and Langerhans cells,” Nat.Rev.Immunol., vol. 2, no. 2, pp. 77–84, 2002.

[9] S. L. Carroll, C. Pasare, and G. M. Barton, “Control of adaptive immunity by pattern recognition receptors,” Immunity, vol. 57, no. 4, pp. 632–648, Apr. 2024, doi: 10.1016/J.IMMUNI.2024.03.014.

[10] P. Stoitzner and N. Romani, “Langerin, the ‘Catcher in the Rye’: an important receptor for pathogens on Langerhans cells,” Eur. J. Immunol., vol. 41, no. 9, pp. 2526–2529, Sep. 2011, doi: 10.1002/EJI.201141934.

[11] B. E. Clausen and J. M. Kel, “Langerhans cells: critical regulators of skin immunity?,” Immunol. Cell Biol., vol. 88, no. 4, pp. 351–360, May 2010, doi: 10.1038/ICB.2010.40.

[12] L. Furio, I. Briotet, A. Journeaux, H. Billard, and J. Peguet-Navarro, “Human langerhans cells are more efficient than CD14(-)CD1c(+) dermal dendritic cells at priming naive CD4(+) T cells,” J.Invest.Dermatol., vol. 130, no. 1523–1747 (Electronic), pp. 1345–1354, 2010.

[13] A. E. Morelli et al., “CD4+ T cell responses elicited by different subsets of human skin migratory dendritic cells,” J. Immunol., vol. 175, no. 0022–1767 (Print), pp. 7905–7915, 2005.

[14] P. Stoitzner et al., “Langerhans cells cross-present antigen derived from skin,” Proc.Natl.Acad.Sci.U.S.A, vol. 103, pp. 7783–7788, 2006.

[15] C. Levin et al., “Critical Role for Skin-Derived Migratory DCs and Langerhans Cells in TFH and GC Responses after Intradermal Immunization,” J. Invest. Dermatol., vol. 137, no. 9, pp. 1905–1913, Sep. 2017, doi: 10.1016/J.JID.2017.04.016/ATTACHMENT/589EE89E-C8C0-4BBF-B164-AC47D4FFCCA2/MMC1.PDF.

[16] C. Yao et al., “Skin dendritic cells induce follicular helper T cells and protective humoral immune responses,” J. Allergy Clin. Immunol., vol. 136, no. 5, pp. 1387–1397.e7, Nov. 2015, doi: 10.1016/J.JACI.2015.04.001.

[17] T. Vardam and N. Anandasabapathy, “Langerhans Cells Orchestrate T FH-Dependent Humoral Immunity,” J. Invest. Dermatol., vol. 137, no. 9, pp. 1826–1828, Sep. 2017, doi: 10.1016/J.JID.2017.06.014.

[18] V. Flacher et al., “Epidermal Langerhans cells rapidly capture and present antigens from C-type lectin-targeting antibodies deposited in the dermis,” J.Invest.Dermatol., vol. 130, pp. 755–762, 2010.

[19] L. Bellmann et al., “Targeted delivery of a vaccine protein to Langerhans cells in the human skin via the C-type lectin receptor Langerin,” Eur. J. Immunol., 2021, doi: 10.1002/EJI.202149670.

[20] V. Flacher et al., “Murine Langerin+ dermal dendritic cells prime CD8+ T cells while Langerhans cells induce cross-tolerance,” EMBO Mol. Med., vol. 6, no. 9, pp. 1191–1204, Sep. 2014, doi: 10.15252/EMMM.201303283.

[21] V. Dioszeghy et al., “Antigen Uptake by Langerhans Cells Is Required for the Induction of Regulatory T Cells and the Acquisition of Tolerance During Epicutaneous Immunotherapy in OVA-Sensitized Mice,” Front. Immunol., vol. 9, no. SEP, p. 1951, Sep. 2018, doi: 10.3389/FIMMU.2018.01951.

[22] J. Idoyaga et al., “Specialized role of migratory dendritic cells in peripheral tolerance induction,” J. Clin. Invest., vol. 123, no. 2, pp. 844–854, Feb. 2013, doi: 10.1172/JCI65260.

[23] E. C. Wamhoff et al., “A Specific, Glycomimetic Langerin Ligand for Human Langerhans Cell Targeting,” ACS Cent. Sci., vol. 5, no. 5, pp. 808–820, May 2019, doi: 10.1021/acscentsci.9b00093.

[24] J. Schulze, M. Rentzsch, D. Kim, L. Bellmann, P. Stoitzner, and C. Rademacher, “A Liposomal Platform for Delivery of a Protein Antigen to Langerin-Expressing Cells,” Biochemistry, vol. 58, no. 21, pp. 2576–2580, May 2019, doi: 10.1021/acs.biochem.9b00402.

[25] M. Rentzsch et al., “Specific Protein Antigen Delivery to Human Langerhans Cells in Intact Skin,” Front. Immunol., vol. 12, Oct. 2021, doi: 10.3389/FIMMU.2021.732298.

[26] N. Rahhal, M. Rentzsch, S. Seiser, C. Freystätter, A. Elbe-Bürger, and C. Rademacher, “Targeted delivery of cytotoxic proteins via lipid-based nanoparticles to primary Langerhans cells,” Nanoscale, vol. 17, no. 7, pp. 4038–4046, Feb. 2025, doi: 10.1039/D4NR03638G.

[27] D. H. Kaplan, M. C. Jenison, S. Saeland, W. D. Shlomchik, and M. J. Shlomchik, “Epidermal Langerhans cell-deficient mice develop enhanced contact hypersensitivity,” Immunity, vol. 23, no. 1074–7613 (Print), pp. 611–620, 2005.

[28] M. J. Barnden, J. Allison, W. R. Heath, and F. R. Carbone, “Defective TCR expression in transgenic mice constructed using cDNA-based alpha- and beta-chain genes under the control of heterologous regulatory elements,” Immunol. Cell Biol., vol. 76, no. 1, pp. 34–40, 1998, doi: 10.1046/J.1440-1711.1998.00709.X.

[29] F. W. Shen et al., “Cloning of Ly-5 cDNA,” Proc. Natl. Acad. Sci. U. S. A., vol. 82, no. 21, pp. 7360–7363, 1985, doi: 10.1073/PNAS.82.21.7360.

[30] P. Stoitzner, N. Romani, A. D. McLellan, C. H. Tripp, and S. Ebner, “Isolation of Skin Dendritic Cells from Mouse and Man,” Methods Mol.Biol., vol. 595, no. 1940-6029 (Electronic), pp. 235–248, 2010.

[31] P. Stoitzner, N. Romani, C. Rademacher, H. C. Probst, and K. Mahnke, “Antigen targeting to dendritic cells: Still a place in future immunotherapy?,” Eur. J. Immunol. 2022, vol. 52, pp. 1909–1924, 2022, doi: 10.1002/eji.202149515.

[32] M. Ghattas, G. Dwivedi, M. Lavertu, and M. G. Alameh, “Vaccine Technologies and Platforms for Infectious Diseases: Current Progress, Challenges, and Opportunities,” Vaccines, vol. 9, no. 12, Dec. 2021, doi: 10.3390/VACCINES9121490.

[33] A. Kubo, K. Nagao, M. Yokouchi, H. Sasaki, and M. Amagai, “External antigen uptake by Langerhans cells with reorganization of epidermal tight junction barriers,” J. Exp. Med., vol. 206, pp. 2937–2946, Dec. 2009, doi: 10.1084/JEM.20091527.

[34] V. Dioszeghy et al., “Differences in phenotype, homing properties and suppressive activities of regulatory T cells induced by epicutaneous, oral or sublingual immunotherapy in mice sensitized to peanut,” Cell. Mol. Immunol., vol. 14, no. 9, p. 770, Sep. 2017, doi: 10.1038/CMI.2016.14.

